# Removing unwanted variation between samples in Hi-C experiments

**DOI:** 10.1101/214361

**Authors:** Kipper Fletez-Brant, Yunjiang Qiu, David U. Gorkin, Ming Hu, Kasper D. Hansen

## Abstract

Hi-C data is commonly normalized using single sample processing methods, with focus on comparisons between regions within a given contact map. Here, we aim to compare contact maps across different samples. We demonstrate that unwanted variation, of likely technical origin, is present in Hi-C data with replicates from different individuals, and that properties of this unwanted variation changes across the contact map. We present BNBC, a method for normalization and batch correction of Hi-C data and show that it substantially improves comparisons across samples, including in a QTL analysis as well as differential enrichment across cell types.

## Introduction

The Hi-C assay allows for genome-wide measurements of chromatin interactions between different genomic regions (Lieberman-Aiden et al., 2009; Wit, Laat, 2012; Dekker et al., 2013; Schmitt et al., 2016; Davies et al., 2017). The use of Hi-C has revealed that the genome is organized in structures at different resolutions such as A/B compartments (Lieberman-Aiden et al., 2009), topologically associated domains (TADs) (Dixon et al., 2012; Nora et al., 2012; Sexton et al., 2012) and loops (Rao et al., 2014). Partly because of the high cost of the assay, the role of inter-personal variation in 3D genome structure is largely unexplored, with the exception of our recent work (Gorkin et al., 2019).

In addition to large-scale structures such as TADs and A/B compartments, there is substantial interest in using Hi-C data to measure specific interactions such as those occurring between regulatory elements and their associated promoters. These interactions are represented as individual cells in the Hi-C contact matrix. Such regulatory interactions do not occur at all distances; an example is enhancer-promoter contacts, which are thought to occur primarily within 1 Mb (Vernimmen, Bickmore, 2015). Methods for detecting such interactions include Fit-HiC (Ay et al., 2014) and HiC-DC (Carty et al., 2017); these methods compare specific contact cells to a background distribution.

Variation and noise in a Hi-C experiment can differ between resolutions and between different types of structures. For example, A/B compartments are estimated using an Eigen decomposition of a suitably normalized contact matrix. We have previously (Fortin, Hansen, 2015) found little-to-no differences between A/B compartments estimated using data from a 1 Mb resolution dilution Hi-C experiment (Lieberman-Aiden et al., 2009) and a 1 Kb resolution in-situ Hi-C experiment on the same cell line (Rao et al., 2014). This observation is specific to A/B compartments; the two experiments differ dramatically in terms of resolution and ability to estimate many other types of structures including TADs and loops.

Hi-C data, like all types of genomic data, suffers from systematic noise and bias. To address this, a number of within-sample normalization methods have been developed. Some of these methods explicitly model sources of unwanted variation, such as GC content of interaction loci, fragment length, mappability and copy number (Yaffe, Tanay, 2011; Hu et al., 2012; Vidal et al., 2018). Other methods are agnostic to sources of bias and attempts to balance the marginal distribution of contacts (Imakaev et al., 2012; Knight, Ruiz, 2013; Rao et al., 2014; Yan et al., 2017). A comparison of some of these methods found high correlation between their correction factors (Rao et al., 2014).

When comparing genomic data *between* samples, variation can arise from numerous sources that do not reflect the biology of interest including sample procurement, sample storage, library preparation, and sequencing. We refer to these sources of variation as “unwanted” here, because they obscure the underlying biology that is of interest when performing a between-sample comparison. It is critical to correct for this unwanted variation in analysis (Leek, Scharpf, et al., 2010). A number of tools and extensions have been successful at this, particularly for analysis of gene expression data (Leek, Storey, 2007; Leek, Storey, 2008; Gagnon-Bartsch, Speed, 2012; Johnson et al., 2007; Stegle et al., 2010; Leek, 2014; Risso et al., 2014). Most existing normalization methods for Hi-C data are single sample methods, focused on comparisons between different loci in the genome.

Three existing methods have considered between-sample normalization in the context of a differential comparison (ATL Lun, Smyth, 2015; Stansfield, Cresswell, Vladimirov, et al., 2018; Stansfield, Cresswell, Dozmorov, 2019), all can be viewed as an adaption of the idea of loess normalization from gene expression microarrays (YH Yang et al., 2002). In these methods, the estimated fold-change between conditions are modeled using a loess smoother as a function of either average contact strength (ATL Lun, Smyth, 2015) or distance between loci (Stansfield, Cresswell, Vladimirov, et al., 2018; Stansfield, Cresswell, Dozmorov, 2019). Using the loess estimates, the data are corrected so there is no effect of the covariate on the fold-change.

## Results

### High-quality Hi-C experiments on different individuals

To investigate the variation between Hi-C data generated from individuals with different genetics, we use existing dilution Hi-C data from lymphoblastoid cell lines generated from 8 different individuals (including 2 trios) from the HapMap project (International HapMap Con0sortium, 2003) (Table 1). The individuals cover 3 populations (Yoruba, Han Chinese and Puerto Rico). For each individual, data was generated from two cultures of the same cell line grown separately for at least 2 passages, and more than 500 million mapped reads were generated for each individual (Table 2); at least 250 million reads for each growth replicate. The reads were summarized at a resolution of 40 kb.

**Table 1.** Sample information.

| Sample | Replicate | Ethnicity | Sex | Family | Role | Batch | Prep date | SCC |
| --- | --- | --- | --- | --- | --- | --- | --- | --- |
| HG00512 | 1 | CHS | M | 2 | Father | B1 | 3/4/15 | 0.923 |
| HG00512 | 2 | CHS | M | 2 | Father | B2 | 5/28/15 |  |
| HG00513 | 1 | CHS | F | 2 | Mother | B1 | 3/4/15 | 0.961 |
| HG00513 | 2 | CHS | F | 2 | Mother | B2 | 5/28/15 |  |
| HG00514 | 1 | CHS | F | 2 | Child | B1 | 3/4/15 | 0.967 |
| HG00514 | 2 | CHS | F | 2 | Child | B2 | 5/28/15 |  |
| HG00731 | 1 | PUR | M | 3 | Father | B1 | 3/4/15 | 0.963 |
| HG00731 | 2 | PUR | M | 3 | Father | B2 | 5/28/15 |  |
| HG00732 | 1 | PUR | F | 3 | Mother | B1 | 3/4/15 | 0.956 |
| HG00732 | 2 | PUR | F | 3 | Mother | B2 | 5/28/15 |  |
| HG00733 | 1 | PUR | F | 3 | Child | B1 | 3/4/15 | 0.971 |
| HG00733 | 2 | PUR | F | 3 | Child | B2 | 5/28/15 |  |
| GM19238 | 1 | YRI | F | 1 | Mother | B3 | 9/26/14 | 0.973 |
| GM19238 | 2 | YRI | F | 1 | Mother | B3 | 9/26/14 |  |
| GM19239 | 2 | YRI | M | 1 | Father | B3 | 9/26/14 | N/A |

**Table 2.** Mapping statistics.

| Sample | Replicate | Total Reads | Cis | Cis (Long) | Trans |
| --- | --- | --- | --- | --- | --- |
| GM19238 | 1 | 545,759,860 | 302,092,644 | 230,613,702 | 243,667,216 |
| GM19238 | 2 | 314,967,258 | 185,913,678 | 145,343,000 | 129,053,580 |
| GM19239 | 2 | 553,838,876 | 367,216,970 | 287,593,654 | 186,621,906 |
| HG00512 | 1 | 311,906,326 | 139,566,774 | 94,984,622 | 172,339,552 |
| HG00512 | 2 | 270,228,888 | 152,292,628 | 114,914,390 | 117,936,260 |
| HG00513 | 1 | 371,772,886 | 174,783,850 | 125,704,946 | 196,989,036 |
| HG00513 | 2 | 277,954,128 | 161,423,552 | 122,711,298 | 116,530,576 |
| HG00514 | 1 | 354,765,444 | 210,846,240 | 103,777,676 | 143,919,204 |
| HG00514 | 2 | 266,032,734 | 177,325,378 | 100,665,340 | 88,707,356 |
| HG00731 | 1 | 324,496,352 | 173,380,026 | 105,098,564 | 151,116,326 |
| HG00731 | 2 | 266,661,686 | 151,763,346 | 99,399,932 | 114,898,340 |
| HG00732 | 1 | 419,151,786 | 237,460,978 | 117,332,062 | 181,690,808 |
| HG00732 | 2 | 291,561,824 | 176,279,418 | 99,373,490 | 115,282,406 |
| HG00733 | 1 | 356,662,684 | 185,558,600 | 112,732,352 | 171,104,084 |
| HG00733 | 2 | 293,167,014 | 178,562,640 | 100,250,886 | 114,604,374 |

Quality control using recently developed guidelines (Yardımcı et al., 2019) suggests that our data is of high quality. In support of this conclusion, we used HiCRep to compute stratum adjusted correlation coefficients (SCCs) between replicates of the same cell line (T Yang et al., 2017). This shows a minimal between-growth-replicate SCC of 0.92 with a mean of 0.96, comfortably exceeding the values recommended by Yardımcı et al. (2019).

In our experimental setup, replicate 1 of GM12239 was prepared in a batch separate from all other samples. For this reason, it is hard to assess the variation within and between batch and this replicate is not included in our analysis.

### Experimental design and replication

We use lymphoblastoid cell lines from the HapMap project (International HapMap Consortium, 2003), because these cell lines have been a widely used model system to study inter-individual variation and genetic mechanisms in numerous molecular phenotypes including gene expression, chromatin accessibility, histone modification, and DNA methylation (Stranger et al., 2007; Pickrell et al., 2010; Montgomery et al., 2010; Degner et al., 2012; Kasowski et al., 2013; McVicker et al., 2013; Kilpinen et al., 2013; Bell et al., 2011). It has been established that phenotypic differences, which are unlikely to be explained by genetics, exists between lymphoblas-toid cell lines from different HapMap populations (Stark et al., 2010; Choy et al., 2008; Stranger et al., 2007). These differences might be related to cell line creation and division (Stark et al., 2010). In our experimental design, experimental batch (library preparation) is partly confounded by cell line population (Table 1), because batch B3 consists solely of samples from the Yoruban population wheres batch B1 and B2 contain one growth replicate each from the samples from the Han Chinese and Puerto Rican populations. In addition, batch B1 and B2 were prepared closer together in time (within 3 months) compared to batch B3 (6 months earlier).

The literature on Hi-C data frequently refers to “biological replicates”, but the definition of this term varies. For example, the ENCODE Terms and Definitions (https://www.encodeproject.org/data-standards/terms/) defines a biological replicate as the same experiment performed on different biosamples, an example is different growths of the same cell line. In contrast, Rao et al. (2014) defines biological replicates to be cells which were not cross-linked together; this is looser than the ENCODE definition. In the literature on population level variation in genomic measurements, biological replicates usually refers to replicates from distinct individuals such as different people or different mice. To avoid confusion in the present manuscript, we will use the term “individual replicate” to refer to a replicate experiment performed on lymphoblastoid cells lines created from two distinct individuals. And we will use the term “growth replicate” to refer to a replicate experiment on a different growth of the same cell line – this is what is commonly referred to as a “biological replicate” in the Hi-C literature.

### Unwanted variation in Hi-C data varies between distance stratum

It is well described that a Hi-C contact map exhibits an exponential decay in signal as the distance between loci increases (Lieberman-Aiden et al., 2009). When we quantify this behavior across growth and individual replicates, we observe substantial variation in the decay rate from sample to sample (Figure 1a). The data presented in Figure 1a is unnormalized, but it strongly suggests systematic unwanted variation between experimental batches, especially batch B3 (Yorubeans) are different from the other two batches. We note that nothing from our quality control analysis suggests that samples from batch B3 have either lower or higher quality than the rest.

**Figure 1.**
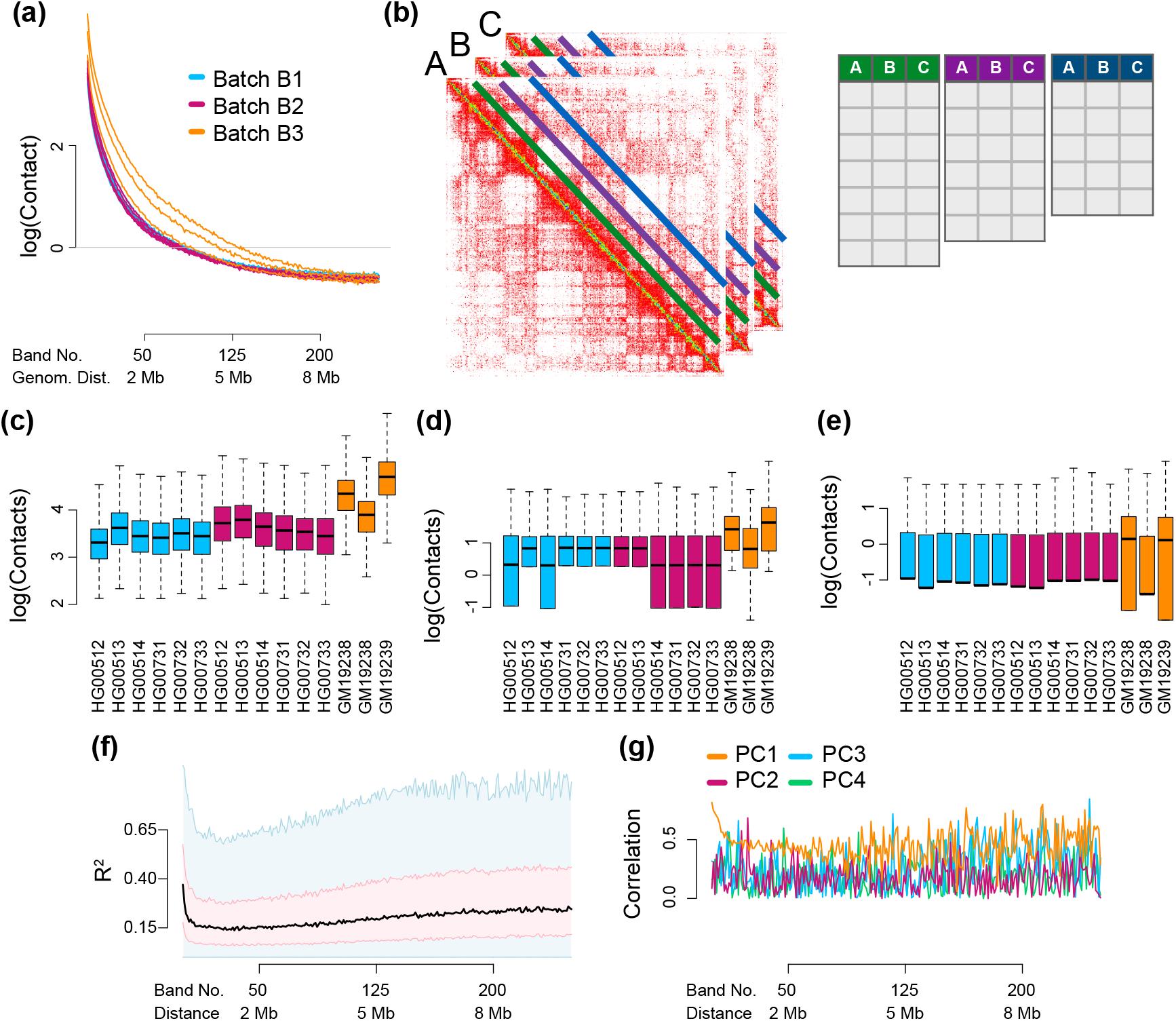
Substantial between-sample variation in unnormalized Hi-C data. We display Hi-C data from chromosome 14 from 8 different individuals, 7 of which have 2 technical replicates, processed in 3 batches. The data has not been normalized apart from correction for library size using the log counts per million transformation; data is on a logarithmic scale. **(a)** Mean contact as a function of distance. Each sample is a separate curve. **(b)** Band transformation of a collection of Hi-C contact maps. **(c)-(e)** Boxplots of the marginal distribution of contacts across samples, for loci separated by **(c)** 40 kb (band 2), **(d)** 2 Mb (band 50) and **(e)** 8 Mb (band 200). **(f)** The percentage of variation explained (*R*^2^) in a linear mixed effect model with library preparation as explanatory variable. The plot is a smoothed boxplot (Methods) with the black line depicting the median, the red shape depicting the 25%- and 75%-quantiles and the blue shape depicting 1.5 times the interquartile range. **(g)** The Spearman correlation of the library preparation factor with each of the first 4 principal components of each band matrix.

In molecular profiling, we frequently observe substantial technical variation in the data. This variation is often associated with experimental batch and has been termed “batch effects” (Leek, Scharpf, et al., 2010). Later, Gagnon-Bartsch, Speed (2012) introduced the more general term “unwanted variation”. What is considered unwanted variation is study specific, and can include stochastic variation, technical variation at the level of sample collection, technical variation at the level of library preparation, and also biological variation of no interest. As an example of the later, in molecular profiling of tissues (but not cell lines as studied here), variation in cell type composition is frequently considered unwanted, but is sometimes the subject of interest. We refer to all of these sources of variation which can obscure the biological differences of interest, as “unwanted variation”.

To assess unwanted variation beyond changes in the mean, we represented our data as a set of matrices indexed by genomic distance (Figure 1b). Each matrix contains all contacts between loci at a fixed genomic distance for all samples (Methods). We call this a band transformation, since these contacts form diagonal bands in the original Hi-C contact matrices (a band in our usage is sometimes described as a matrix diagonal; it is a “line” parallel to the main matrix diagonal). Figure 1c-e depicts the distributions for three selected bands at close (40 kb), medium (2 Mb) and long (8 Mb) ranges. These distributions suugests that band-specific variation is also systematically different between batches.

To quantify the impact of unwanted variation on our Hi-C data, we first asked, for each entry in the contact matrix, how much variation is explained by the experimental batch factor? A factor is a statistical term to describe a variable with a discrete number of states, which has an associated code (here B1, B2 and B3, see Table 1). We measure the amount of explained variation using *R*^2^ from a linear mixed effects model with a random effect to model the increased correlation between growth replicates (Methods). We observe an association between explained variation and distance between loci (Figure 1)f, with an average *R*^2^ value of 0.23. This means that 23% of the between-sample variation in the individual entries in the contact matrix is explained by experimental batch, which is partly confounded with population (explored further below). This shows that the effect of the experimental batch factor changes with distance and is substantial.

To further explore the effect of batch, we performed principal component analysis on each of the band matrices and computed Spearman correlation between each of the first four principal components and the batch indicator (Figure 1). This is a common technique to assess if the major sources of variation in a matrix is associated with a known covariate, here the experimental batch factor. This supports the conclusion of our *R*^2^ analysis and emphasizes the dynamic nature of the association between variability and the experimental batch factor.

### ICE and observed-expected normalization between samples

A number of normalization methods have been developed for Hi-C data, many with the explicit purpose of removing bias along the genome (see the Introduction). Many of these methods do not explicitly contain even a scaling normalization between samples. A popular method is ICE (Imakaev et al., 2012).

Observed-expected normalization was introduced by Lieberman-Aiden et al. (2009); it consists of dividing all contact cells in a given band by the mean contact across the band. This is an example of scale normalization, and was introduced as a *within-sample* normal-ization technique. In light of the differences in decay rates across samples (Figure 1a), it is natural to force the decay rates to be the same. Observed-expected normalization is an easy approach to this, since it removes the decay and hence forces different samples to have the same (non-existing) decay rate. To keep the fast decay rate in the data, we suggest multiplying the band matrices by the average decay rate (Methods). This is a natural adaptation of observed-expected normalization to a *between-samples* approach. We combine observed-expected between-sample normalization with ICE, and we refer to this combined method as ICE-OE.

Using ICE-OE leads to a substantial improvement (Figure 2). Per design, there is no between sample variation in the contact decay rate. Boxplots of the contact distribution for selected bands still show sample-specific variance. More important, we no longer observe any dependence of *R*^2^ on band, and the average *R*^2^ is at the level of the smallest *R*^2^ for unnormalized data (ie. 0.15). While the *R*^2^ is smaller than for unnormalized data alone, we note that, for each distance band, 25% of the contact cells still have an *R*^2^ of 0.3 or greater. Likewise we observe improvement in the correlation between principal components and the batch factor. With these assessments, ICE-OE appears to have addressed many of the major deficiencies associated with unnormalized data.

**Figure 2.**
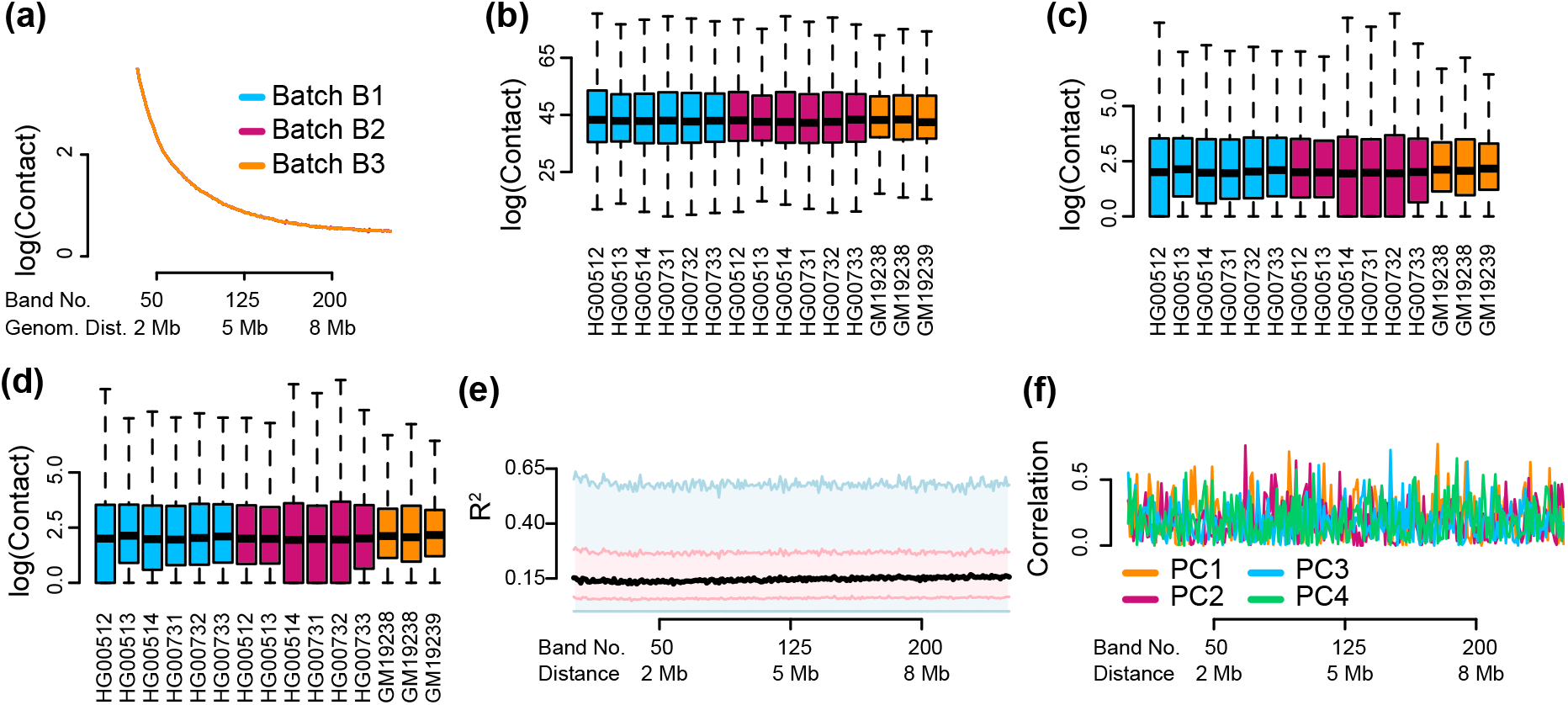
Unwanted variation in Hi-C data normalized by ICE-OE. Hi-C data is normalized by ICE followed by observed-expected normalization across samples (ICE-OE). **(a)** Mean contact as a function of distance. **(b)-(d)** Boxplots of the marginal distribution of contacts across samples, for loci separated by **(b)** 40 kb (band 2), **(c)** 2 Mb (band 50) and **(d)** 8 Mb (band 200). **(e)** The percentage of variation explained (*R*^2^) in a linear mixed effect model with library preparation as explanatory variable. **(f)** The Spearman correlation of the library preparation factor with each of the first 4 principal components of each band matrix.

There is some difference between existing methods for normalization of Hi-C data matrices. Figure 3 show the *R*^2^ smoothed boxplot for data normalized using HiCNorm (Hu et al., 2012) (but without the observed-expected modification we use for ICE-OE). We observe a substantial dependence of *R*^2^ on distance.

**Figure 3.**
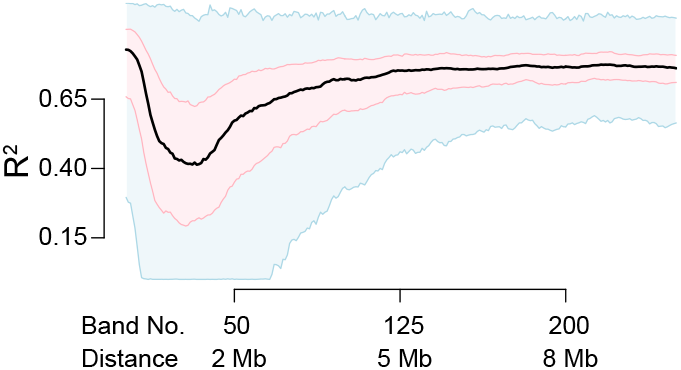
Unwanted variation and HiCNorm. Like Figures 1f and 2e, but for data normalized using HiCNorm.

### The separate impact of library preparation and cell line creation

It is interesting to consider the source of the unwanted variation. To investigate this, we restricted our analysis to samples from the Han Chinese and Puerto Rican populations since – for these samples – each growth replicate was prepared twice in two different batches (Table 1). This results in a balanced experiment, making it easy to separate the contribution of these two factors.

Using the *R*^2^ approach described above, we can compute *R*^2^ for a model only including population (cell line creation) and a model only including library preparation. These two sets of explained variation are comparable because the data is unchanged and the explanatory variable has the same dimension with the same number of replicates assigned to each level. We can also compute *R*^2^ for a model containing both factors; mathematically this is guaranteed to be higher than *R*^2^ for either factor separately. We do this both for unnormalized data and for data processed using ICE-OE. In Figure 4, we observe that library preparation explains slightly higher variation compared to population. For the unnormalized data, there is substantial variation of the effect of library preparation in different bands, which goes away after the observed-expected normalization. We also observe that the two factors appear to combine independently, in that including both factors in a model raises *R*^2^ substantially above either of the two factors alone. We conclude that library preparation explains at least as much variation as cell line creation, and possibly more.

**Figure 4.**
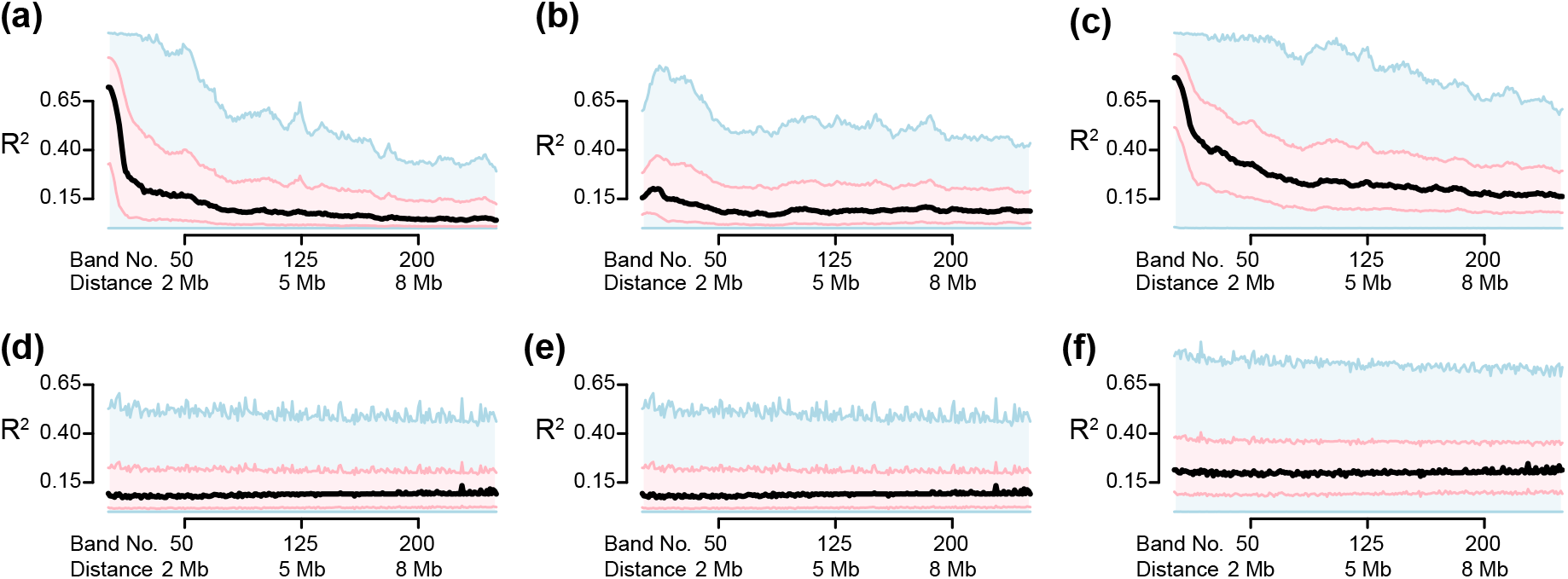
The impact of library preparation and cell line creation on unwanted variation. We use Hi-C data from 6 individuals from 2 different populations (cell line creation). Each individual has two library preparation replicates performed at two time points, allowing us to separate the effect of population (cell line creation) and library preparation. The effect of these two variables are quantified through the percentage of variance explained (*R*^2^) in different models where either population (cell line creation) or library preparation or both are included. **(a)** Unnormalized Hi-C data, model only includes library preparation. **(b)** Unnormalized Hi-C data, model only including population (cell line creation). **(c)** Unnormalized Hi-C data, model including both library preparation and population (cell line creation). **(d)-(f)** Like (a)-(c) but for data normalized using ICE-OE.

Previously, we noted that batch B3 appeared to be more variable than batch B1 and B2. However, here we restrict our analysis to only these later batches and observe a similar percentage of variance explained by batch. This shows that our results above are not just driven by batch B3.

### ICE-OE is unsuitable for genetic mapping

An interesting biological question, which can only be addressed with data on individual replicates, is the association between genetic variation and 3D structure. This question can be asked for any type of 3D structure including TADs and loops. Here we focus on variation in individual contact cells, which is interesting because of the relationship between regulatory elements and the genes they regulate. Specifically we are interested in performing a quantitative trait loci (QTL) mapping for each contact cell. A QTL mapping is simply asking, for each contact cell, whether there is an association with a nearby single nucleotide variant (SNP). An advantage of QTL mapping is that we have well-established quality control procedures which can help reveal whether a data matrix has been properly normalized, which are the focus of the present manuscript. We recently described a complete investigation of the genetic contribution to 3D genome structure (Gorkin et al., 2019).

For our QTL mapping we consider all contact cells representing loci separated by less than 1 Mb. For testing against a given contact cell, we require a candidate SNP to be present in one of the two anchor bins for that contact cell (Figure 5a, Methods). We furthermore require that all genotypes are represented in our samples (Methods). These requirements yield a total of 22,541 SNPs for 21,017 contact cells on chromosome 22, representing 1,111,407 tests. We use a linear mixed effect model with a random effect on the growth replicate, to model the increased correlation between growth replicates.

**Figure 5.**
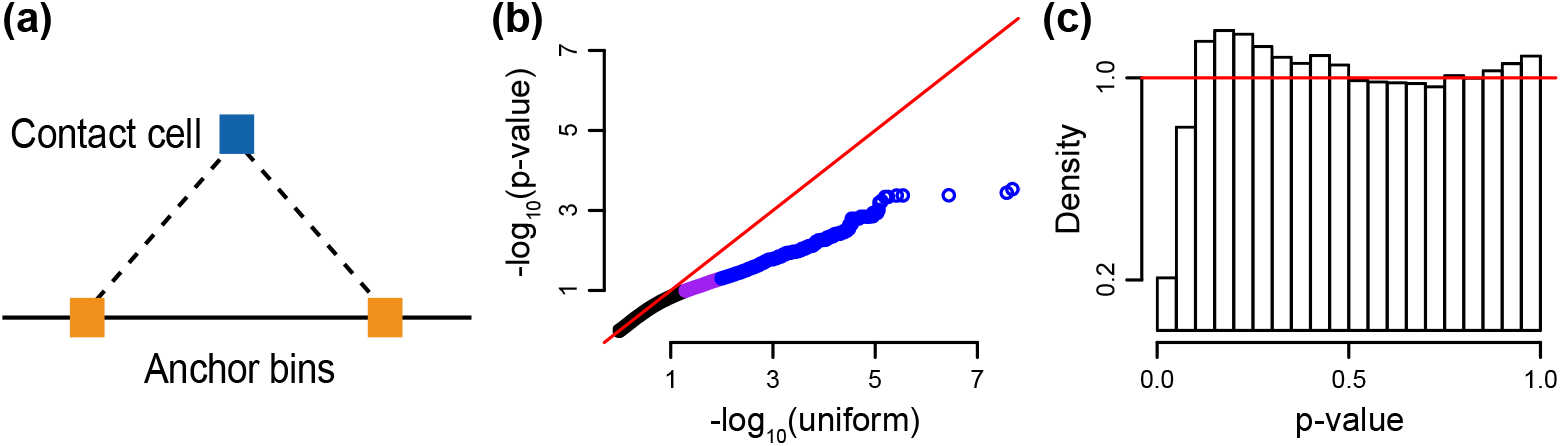
A QTL analysis reveals incorrect normalization. We performed a QTL analysis for association between suitably chosen SNPs on chromosome 22 and contact cells. Data was normalized using either ICE or ICE-OE. **(a)** A diagram of the search procedure. We restricted SNPs to be present in the anchor bins. **(b)** QQ-plots comparing the -log_10_ p-value (x-axis) to log_10_ quantiles from the uniform distribution for data normalized using ICE-OE. Colors: blue the p-value is in the 99*th* percentile or greater, purple if the p-value is in the 95*th* to 99*th* percentiles, and black otherwise. Note that no test has a p-value less than 10^−6^. **(c)** A histogram of the p-values from (a).

In Figure 5 we depict a quantile-quantile plot (QQ-plot) for the (minus logarithmic) p-values for this analysis, as well as histograms of the p-value distribution. We observe that the QQ-plots look unsatisfactory with an unusual discrepancy from expectation (parallel with the *y* = *x* line with a deviation towards the end). Furthermore, the p-value histograms are also strongly deviating from the expected behaviour of being flat with a possible bump near zero. We stress that the lack of small p-values revealed by the histogram is not caused by lack of power due to small sample size; this would result in a flat histogram. We conclude that ICE-OE does not properly normalize the data for a QTL analysis.

### Band-wise normalization and batch correction

To normalize the data and remove unwanted variation for a QTL analysis, we used the band transformation framework. We propose to separately smooth each contact matrix, apply the band transformation, quantile normalize each bandmatrix, followed by using ComBat with a known batch effect factor. We call this approach band-wise normalization and batch correction (BNBC). We next describe our rationale for each step.

We start by following existing work by T Yang et al. (2017) and smooth the sample-specific contact matrices, since doing so results in increased correlation between growth replicates (confirmed by us). We note that the HiC-Rep criteria does not include consideration of biological signal and we caution that such signal could be diminished. For example, in work on normalization of DNA methylation arrays, we found that methods which performs best at reducing technical variation do not necessarily perform best when the assessment is replication of biological signal (Fortin, Labbe, et al., 2014). For these reasons we consider smoothing an optional part of BNBC and our software makes it easy to disable.

We next process each smoothed matrix band, from all samples, one at a time. Chromosomes are processed separately. We perform quantile normalization on each matrix band. Quantile normalization forces the marginal distributions of each sample to be the same, ie. the distributions displayed in Figures 1a,c-e, 6a-d. This reduces inter-sample variability, but operates under an assumption that the genome-wide distribution of contacts at a given distance, is the same across samples. This assumption is in our view uncontroversial for our lymphoblastoid cell lines. Quantile normalization can be disabled in our software.

We then use ComBat (Johnson et al., 2007) to remove the effect of batch in each band matrix separately. ComBat removes the effect of batch on both the location (mean) of a given Hi-C matrix cell’s observations across samples, as well as the scale (variance). In comparison, regressing out the batch factor using a standard linear model would only remove the effect of batch on location (mean). Moreover, ComBat uses Empirical Bayes methods to regularize estimates of batch effects, resulting in more stable estimates, particular in the small-sample setting. In this approach it is important to condition on distance because the exponential decay of the contact matrices would make contact cells from different bands incomparable.

The use of ComBat requires at least 2 replicates in each experimental batch. In our section on the Hi-C experiments we noted that one individual had a replicate prepared in a batch separately from all other samples. Such a replicate cannot be included in the normalization, which is a clear limitation.

BNBC is highly scalable because we only process one matrix band at a time. The largest band – the diagonal – has a number of entries equal to the number of bins in the genome, and this size scales linearly with resolution. A 1kb resolution Hi-C experiment has 3M entries in its diagonal, resulting in a band matrix with 3M rows and columns equal to the number of samples. For further scalability, we process each chromosome separately. While big, this can be processed on a laptop. We provide an implementation supporting the cooler format (Abdennur, Mirny, 2019).

BNBC removes any between sample difference in decay rate and also stabilizes band-specific variances across samples (Figure 6). To assess the impact of BNBC, we again measured the variation explained by the batch factor. We observe a decrease in this quantity compared to ICE-OE, including at the 75%-quantile level (Figures 6, 2). Likewise, we observe a dramatic decrease in correlation between principal components and the batch factor.

**Figure 6.**
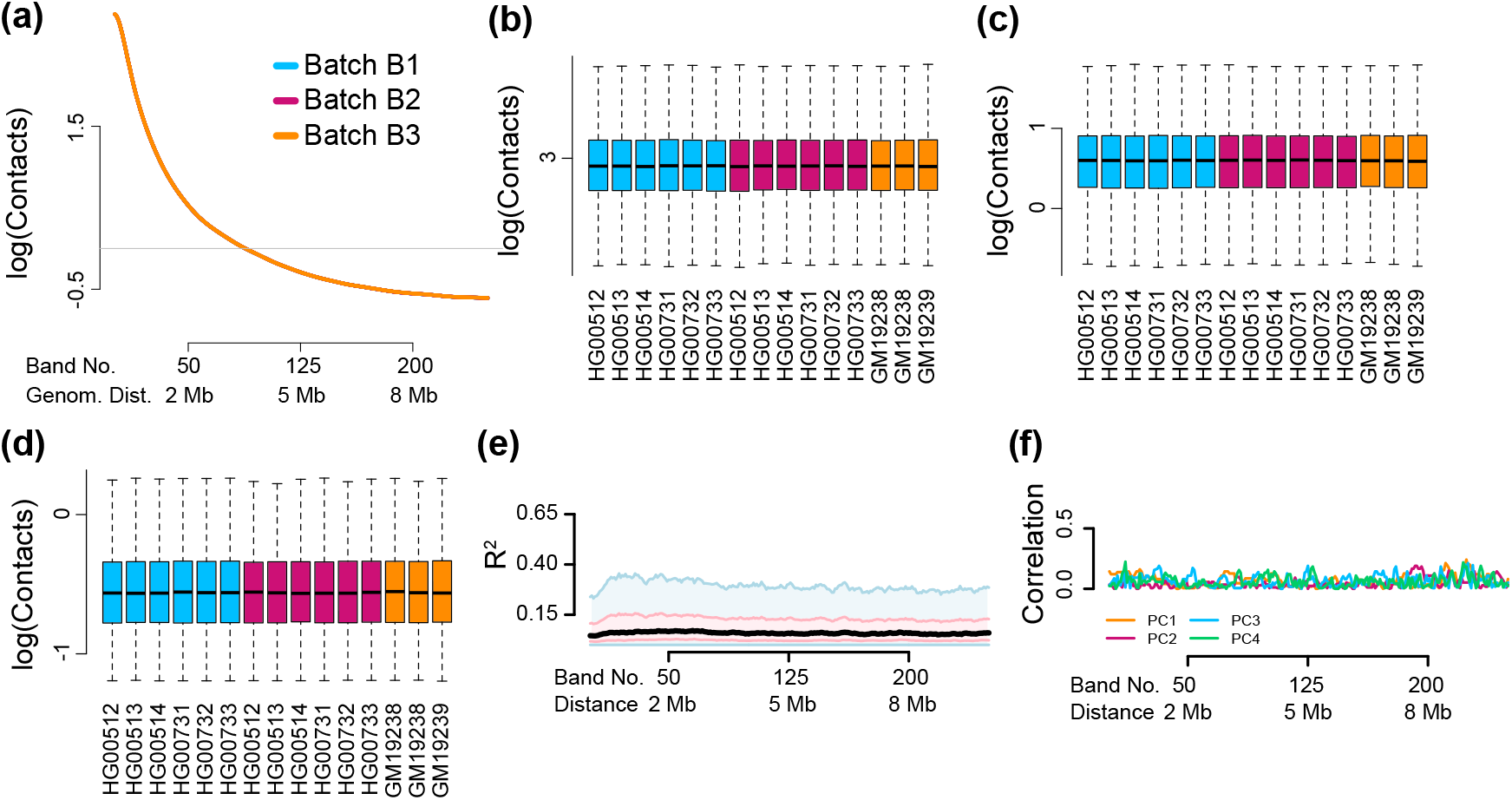
BNBC removes unwanted variation in Hi-C data. As Figure 2 but for data processed using BNBC. **(a)** Mean contact as a function of distance. **(b)-(d)** Boxplots of the marginal distribution of contacts across samples, for loci separated by **(b)** 40 kb (band 2), **(c)** 2 Mb (band 50) and **(d)** 8 Mb (band 200). **(e)** The percentage of variation explained (*R*^2^) in a linear mixed effect model with library preparation as explanatory variable. **(f)** The Spearman correlation of the library preparation factor with each of the first 4 principal components of each band matrix.

While seemingly impressive, we note that the decrease in *R*^2^ and the lack of correlation with principal components are mathematical consequences of the use of regression in ComBat. This is because simply regressing out a factor from each of the entries in a band matrix, ensures that both *R*^2^ for that factor as well as the correlation between factor and each of the principal components of the data matrix is equal to zero. ComBat does more than simply regressing out batch – it uses Empirical Bayes techniques to shrink the parameters and it also models changes in variation – and this explains why the observed *R*^2^ and the correlations are not exactly 0. For this reason, we caution against the use of these evaluation criteria for assessing the performance of BNBC. The assessment of non-regression based techniques, like ICE and ICE-OE, is not impacted by this comment.

We next investigated the impact of BNBC on the contact map. There is little difference between the contact map following ICE normalization and BNBC normlization (Figure 7). The same is true for the associated first Eigenvector, which is commonly used to identify A/B compartments (Figure 7). The correlation between the compartment vectors obtained using BNBC and ICE-OE is 0.959 for GM19238 and 0.957 for HG00513. We conclude that BNBC does not distort gross features of the contact map.

**Figure 7.**
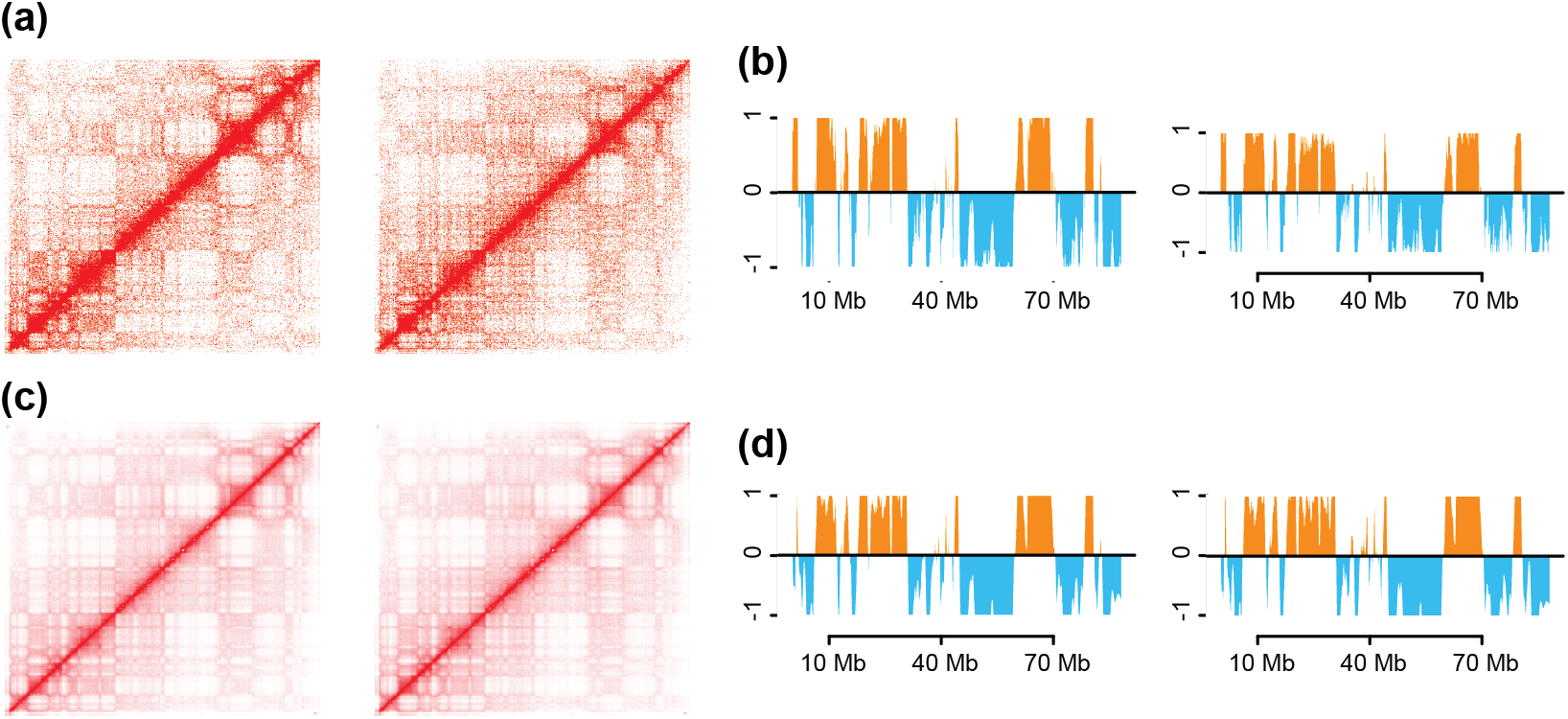
BNBC preserves features of the contact map. Two samples (left: GM19238, right: HG00512) from two different batches are processed using either ICE-OE or BNBC. We display data from chromosome 14. **(a)** Contact maps with data processed using ICE-OE. **(b)** A/B compartments using the 1st principal component of the observed-expected normalized contact matrix with data processed using ICE-OE. **(c)-(d)** Like (a)-(b) but with data processed using BNBC.

We now consider the impact of BNBC on genetic mapping. Using the same measures as described above, we observe a uniform distribution of p-values as well as a much better behaved QQ-plot for the p-values (Figure 8). Multiple observed p-values are less than 10^−6^ (we do more than 1M tests), comfortably exceeding the lowest p-value following ICE or ICE-OE (which is around 10^−4^, Figure 5), suggesting that BNBC not only corrects issues with under-inflation of the test statistic, but also increases power. We conclude that BNBC noticeably improves on ICE and ICE-OE for genetic mapping.

**Figure 8.**
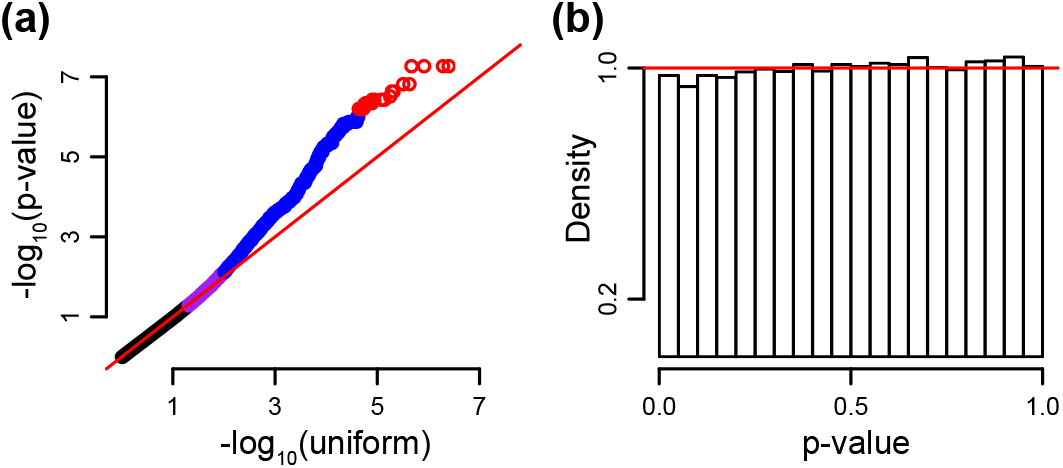
BNBC normalizes data for QTL analysis. Like Figure 5 but for data normalized using BNBC. **(a)** QQ plots comparing the -log_10_ p-value (x-axis) to − log_10_ quantiles from the uniform distribution. Colors: red if the p-value is less than 10^−6^, blue if the p-value is in 99*th* percentile or greater, purple if the p-value is in the 95*th* to 99*th* percentiles, and black otherwise. **(b)** A histogram of the p-values from (a).

### BNBC accommodates sparse contact matrices and substantial coverage variation

To demonstrate the scalability of our approach, and to evaluate BNBC on a separate dataset, we next considered the setting of a sparse, 5k-resolution contact matrix. Specifically, we consider the set of interactions on chr22 from (Greenwald et al., 2019), a study comparing contact propensity between induced pluripotent stem cells (iPSCs) and iPSC-derived cardiomyocytes (CMs), in 7 members of a multi-generational family using in-situ Hi-C. Here we demonstrate that BNBC is applicable to process sparse 5kb matrices and also increases power to detect contact matrix cells which are differentially enriched between cell types.

Hi-C libraries were generated at two different time points (Table 3), with the majority of libraries being generated at one time point (and a few libraries generated at a single different time point). The publicly available processed data is at the (subject, cell type) level and different samples are the result of different pooling strategies. First, some samples were generated by using only a single library and some samples were generated by pooling two libraries. As all libraries were sequenced to approximately the same depth, this creates a substantial fixed difference in library size which – to a first approximation – can be explained by whether the sample was pooled or not (column “Pooled” in Table 3, compared to column “Library Size”). Second, when a sample resulted as a pool of two different libraries, sometimes the two Hi-C libraries were prepared in different batches or not (column “MixBatch” in Table 3). Together, these two variables (“Pooled” and “MixBatch”) creates 3 different levels since there are no samples that mixed batches and were not pooled.

**Table 3.** Greenwald Sample Information.

| Sample | Cell Type | MixBatch | Pooled | Batch | Library Size |
| --- | --- | --- | --- | --- | --- |
| iPSCORE 2.1 | iPSC | no | yes | B1 | 1646980 |
| iPSCORE 2.1 | CM | yes | yes | B2 | 1490690 |
| iPSCORE 2.2 | iPSC | no | yes | B1 | 1505519 |
| iPSCORE 2.2 | CM | no | no | B3 | 703848 |
| iPSCORE 2.3 | iPSC | yes | yes | B2 | 1718901 |
| iPSCORE 2.3 | CM | yes | yes | B2 | 1707640 |
| iPSCORE 2.4 | iPSC | no | no | B3 | 797648 |
| iPSCORE 2.4 | CM | no | yes | B1 | 1454185 |
| iPSCORE 2.6 | iPSC | no | no | B3 | 920916 |
| iPSCORE 2.6 | CM. | no | no | B3 | 695501 |
| iPSCORE 2.7 | iPSC | no | no | B3 | 865444 |
| iPSCORE 2.7 | CM | no | yes | B1 | 1467740 |
| iPSCORE 2.9 | iPSC | yes | yes | B2 | 1928327 |
| iPSCORE 2.9 | CM | no | yes | B1 | 1286331 |

As a first step to investigate unwanted variation, it is helpful to look at the mean contact as a function of distance (Figure 9a). Our preprocessing steps are to apply a modified logCPM transformation that accommodates sparsity to the set of chr22 matrices (Methods), and smooth each contact matrix. This plot reveals evidence of systematic variation between the 3 possible combinations of “Pooled” and “MixBatch”. It may be surprising to see an effect of “Pooled” – this variable is largely describing variation in library size and the logCPM we plot are corrected for library size – but we hypothesize this is caused by the high sparsity of the data where a doubling in library size has a large effect on the sparsity pattern. The effect of “MixBatch” is more in line with the effect we have seen in the analysis of the LCL data (above), although we stress that while we have two library preparation dates, the effect of library preparation date is mitigated by the pooling strategy.

**Figure 9.**
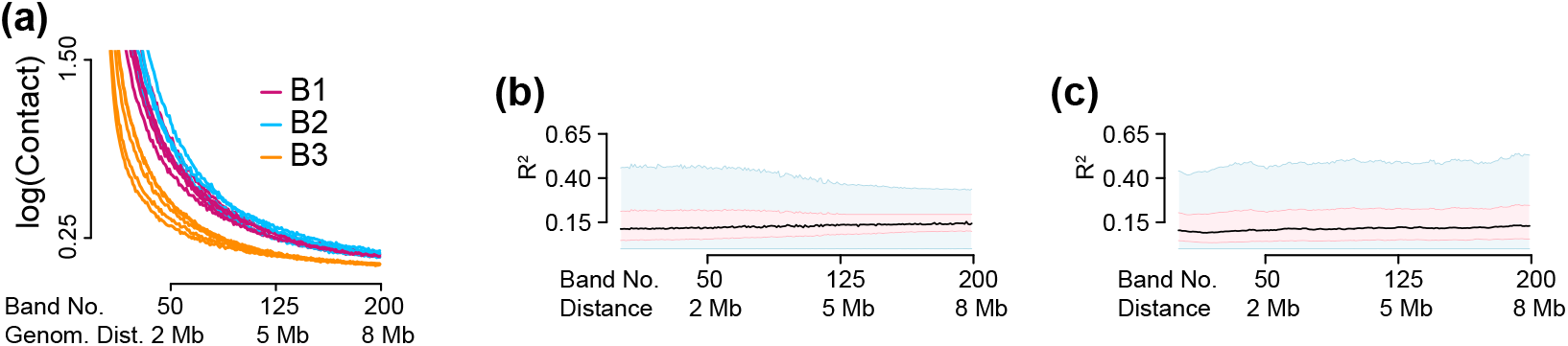
Unwanted Variation in Greenwald et al. Data. **(a)** Mean contact strength as a function of distance in unnnormalized data, stratified by **MixBatch+Pooled** status. **(b)** Partial *R*^2^ as a function of distance, for data processed with ICE-OE. **(c)** As in (b), but for data processed with BNBC.

We next use partial *R*^2^ to compare ICE-OE and BNBC (Figure 9. We need to use partial *R*^2^ in this analysis, because we are asking about the effect of the batch factor in a model which includes a systematic effect of cell type. ICE-OE and BNBC appears to produce data without substantial unwanted variation by this measure, with relatively little difference, except a slight dependence of *R*^2^ on distance for ICE-OE, but with a slight improvement at partial *R*^2^ at long distances compared to BNBC.

### BNBC enhances discovery power while preserving fine-scale detail

To assess BNBC’s discovery power relative to ICE-OE, we conduct a differential contact analysis between cell types, amongst the first 200 matrix bands (see Methods). Examining Figure 10a, it is clear that ICE-OE p-values exhibit unusual behavior inconsistent with the assumption of null p-values following a uniform distribution. Additionally, there is virtually no enrichment of p-values closer to 0, suggesting that ICE-OE is neither calibrated nor powered to detect systematic variation across samples. By contrast, BNBC has clear enrichment of smaller p-values, indicating substantially improved power to detect signal (Figure 10b), with much improved calibration compared to ICE-OE.

**Figure 10.**
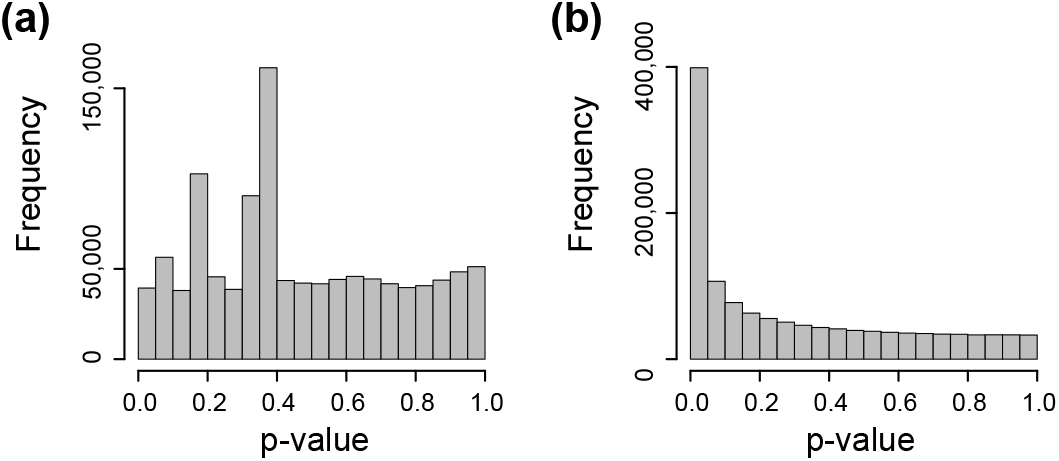
Differential enrichment in the Greenwald data. **(a)** p-value histogram from cell-type differential enrichment analysis using data processed by ICE-OE. **(b)** As (a) but for p-values obtained from BNBC-processed data.

Quantifying the utility of either method, we report the number of hits discovered by either method (Table 4) at a family-wise error rate (FWER) of 0.05 (using a Bonferroni corrected), as well as false discovery rate (FDR) of 0.05, 0.01 and 0.001 (using independent hypothesis weighting, Methods). At no threshold does ICE-OE produce discoveries. Conversely, BNBC is well-powered to make discoveries. Considering stringent thresholds, using Bonferroni to control FWER yields 1,485 significant matrix entries, while we observe 28,354 significant matrix entries at a FDR of 0.001. We used the ICE implementation from Juicer (Durand et al., 2016) which has its own filtering steps and for that reason we perform fewer tests for ICE-OE compared to BNBC. When we restrict BNBC to the entries kept by Juicer, we get the same behaviour.

**Table 4.** Number of significant tests for differential enrichment.

| Method | Number of Tests | Bonferroni | IHW | IHW | IHW |
| --- | --- | --- | --- | --- | --- |
|  |  | FWER < 0.05 | FDR < 0.05 | FDR < 0.01 | FDR < 0.001 |
| ICE-OE | 1,099,648 | 0 | 0 | 0 | 0 |
| BNBC | 1,269,066 | 1,485 | 229,076 | 104,507 | 28,254 |
| BNBC on ICE entries | 1,099,648 | 1,571 | 215,792 | 100,342 | 27,533 |

Finally, to establish that BNBC does in fact preserve local contact map features, we visualize a 3 MB region (at 5 kb resolution) of chr22 (chr22:27000000-30000000; Figure 11) using HiGlass (Kerpedjiev et al., 2018). To highlight finer detail (while allowing for clear visualization of some larger features) we zoom into a 2MB window (chr22:2800000-30000000), making the individual matrix cell contributions evident. While the results of ICE-OE and BNBC are technically on different scales, it is clear that both coarse and fine details are preserved by BNBC as compared to ICE-OE.

**Figure 11.**
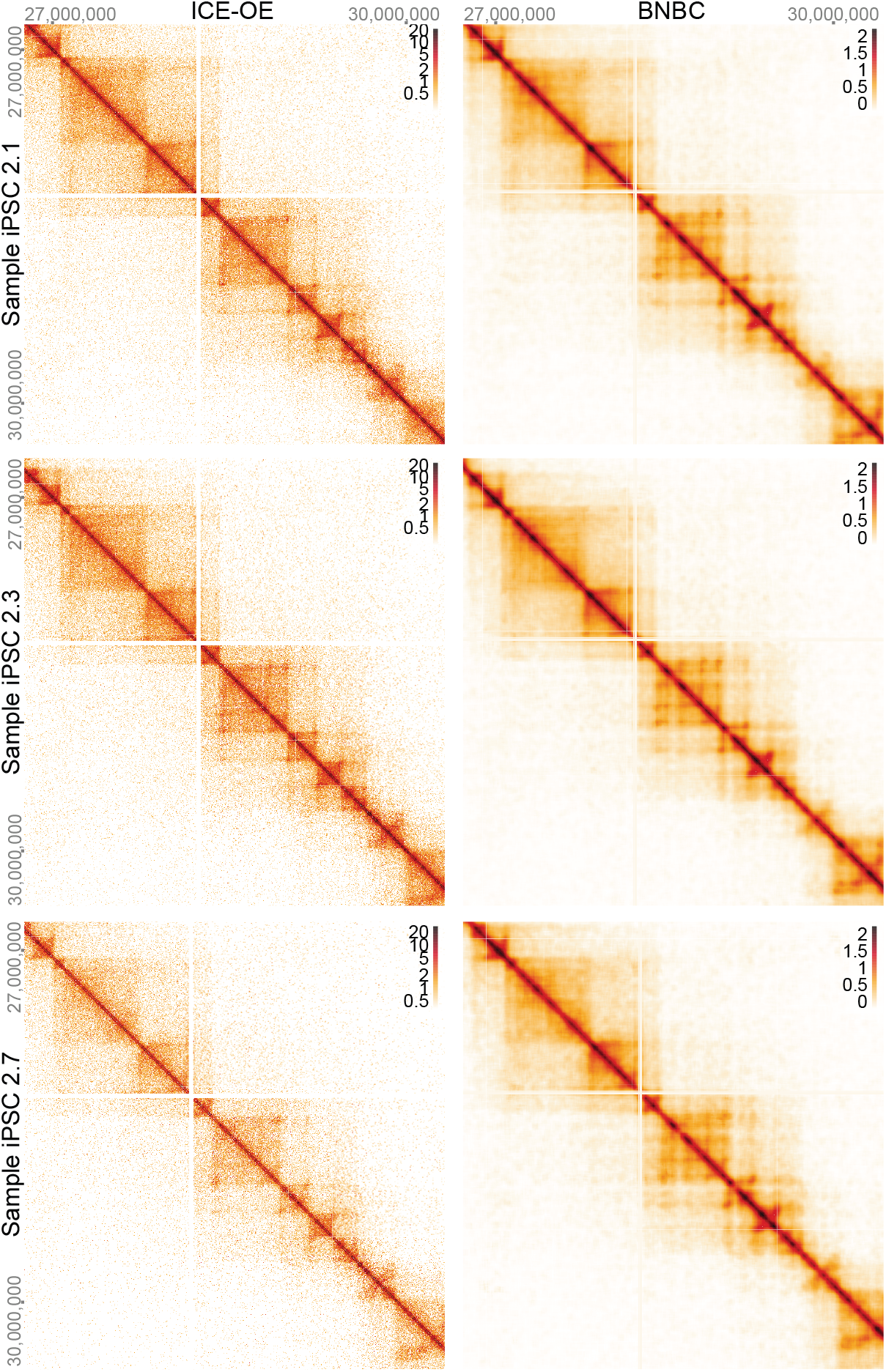
Stucture of Hi-C data following BNBC and ICE-OE. Data is 3 samples from Greenwald et al. (2019). We display 3MB from chr22 at a 5 kb resolution. Data in the left column has been processed by ICE-OE, data in the right column by BNBC.

## Discussion

Here, we have characterized unwanted variation present in Hi-C contact maps and have developed a correction method named band-wise normalization and batch correction (BNBC). We show the existence of unwanted variation in Hi-C data and show that on average, experimental batch explains 32% of the between-sample variation in contact cells for ICE normalized data, in the 40kb dilution Hi-C experiment analyzed here. We show unwanted variation exhibits a distance-dependent effect, in addition to known distance-based features of Hi-C contact maps. A simple combination of ICE and observed-expected normalization adapted to a between-sample normalization method corrects several of these deficiencies; we call this approach ICE-OE. We show that both ICE and ICE-OE has serious deficiencies when used for genetic mapping.

We present BNBC, a modular approach where we combine band transformation with existing tools for normalization and removal of unwanted variation for between-sample comparisons. This is not a method suitable if the intention is to pool data from different replicates into a single contact matrix. We show that BNBC performs well in reducing the impact of unwanted variation while still preserving important 3D features, such as the structure of the contact map and A/B compartments. Data processed using BNBC shows dramatic improvement when used for genetic mapping.

A limitation of our method is the requirement for an explicit batch factor, caused by the use of ComBat. For gene expression analysis, models based on factor analysis such as RUV (Gagnon-Bartsch, Speed, 2012; Risso et al., 2014) or SVA (Leek, Storey, 2007; Leek, Storey, 2008; Leek, 2014) do not have this limitation and has shown outstanding performance. It will be useful to adapt such approaches to Hi-C data analysis. As always, it is important that the experimental setup (a possible batch factor) does not confound the comparison of interest.

We apply ComBat separately to each band, which allows for runtime to increase only linearly in the number of bands to be normalized. A natural extension to ComBat would be a hierarchical model across bands; such a model would allow us to borrow information across bands. However, such a model would not have the same computational scaling properties as the setup we propose. We leave the development of such a model to future work.

In terms of alternative methods, our approach is most readily compared with multiHiCcompare (Stansfield, Cresswell, Dozmorov, 2019), another multi-sample normalization method. The starting point for multiHiC-compare is a pairwise comparison between two samples. Loess is used to model the mean contact difference between the two datasets as a function of distance (band). To get a multi-sample method, all pairwise comparisons are made until convergence. Based on analysis for DNA microarrays, we would expect this method to perform similarly to quantile normalization in each band. In addition to this step, BNBC removes the effect of a batch variable, ie. a specific systematic difference between samples. Such a step is not performed for multiHiCcompare and – based on experience from gene expression analysis – implies that a potential batch effect is not removed.

As a by-product of both ICE-OE and BNBC we force the decay in contact probabilities over distance to be the same across samples. Haarhuis et al. (2017) reports that knockdown of WAPL in HAP1 cells results in changes in the contact decay probability. However, we show in our work that replicates can have quite different decay rates, which suggests that one should be careful before making claims about changes in decay rate. If decay rates are different across samples, forcing them to be similar will remove some biological signal and care should be taken with analysis.

Our approach does not employ the matrix balancing normalization, which is standard in Hi-C analysis. Matrix balancing is used to resolve confounders between different matrix entries from the same sample, such as GC content. Removing such confounders is important when comparing across matrix entries within a matrix from the same sample. In our work we are comparing individual matrix entries across replicates. Because we never compare different matrix entries to each other, this type of comparison is conditional on factors which depends on genomic location (such as GC content). This explains the seeming paradox of matrix balancing being widely used in single-sample Hi-C analysis, yet we found it to be insufficient for comparisons of individual matrix entries across samples.

We have found little reason to process the entire genome at once and have instead opted to processed each chromosome separately. In addition, an anonymous reviewer has pointed out the potential for issue near chromosome arms where there is uncertainty of the length of the centromere. For this reason, it may be desirable to process each arm separately. We note that our evaluation data on chromosome 22 is on a single arm.

We remove genomic bins with poor mapability, low GC content and short fragment lengths. An alternative approach is simply to remove bins with little coverage.

We emphasize that our analysis of unwanted variation is about variation at the level of individual contact cells. The amount of unwanted variation can depend on the type of structure of interest such as TADs or loops. It is not self-evident that using BNBC on the contact matrix is suitable for normalizing TADs or loops for comparisons across samples; this question is not examined in our work.

In summary, proper normalization and correction for unwanted variation will be critical for comparing Hi-C contact maps between different samples.

## Methods

### Data Generation

#### Hi-C experiments

Lymphoblast Hi-C data analyzed were generated by the dilution Hi-C method using HindIII (Lieberman-Aiden et al., 2009) on 9 lymphoblastoid cell lines derived from the 1000 Genomes project (Table 1). Data are publicly available through 1000 genomes (Chaisson et al., 2019) as well as through the 4D Nucleome data portal (https://data.4dnucleome.org; accessions 4DNESYUYFD6H, 4DNESVKLYDOH, 4DNESHGL976U, 4DNESJ1VX52C, 4DNESI2UKI7P, 4DNESTAPSPUC, 4DNES4GSP9S4, 4DNESJIYRA44, 4DNESE3ICNE1). Hi-C contact matrices were generated by tiling the genome into 40kb bins and counting the number of interactions between bins. We refer to these as raw contact matrices.

#### Hi-C read alignment and contact matrices

Reads were aligned to hg19 reference genome using bwa-mem (Li, 2013). Read ends were aligned independently as paired-end model in BWA cannot handle the long insert size of Hi-C reads. Aligned reads were further filtered to keep only the 5’ alignment. Read pairs were then manually paired. Read pairs with low mapping quality (MAPQ¡10) were discarded, and PCR duplicates were removed using Picard tools 1.131 http://broadinstitute.github.io/picard. To construct the contact matrices, Hi-C read pairs were assigned to predefined 40Kb genomic bins. Bins with low mapping quality (*<* 0.8), low GC content (*<* 0.3), and low fragment length (*<* 10% of the bin size) were discarded.

### Band Matrices

To make comparisons across individuals, we form band matrices, which are matrices whose columns are all matrix band *i* from each sample. A matrix band is a collection of entries in a contact matrix between two loci at a fixed distance. Formally, band *i* is the collection of *j*, *k* entries with |*j* − *k*| + 1 = *i*.

### Log counts per million transformation

We use the logCPM (log counts per million) transformation previous described (Law et al., 2014). Specifically, for a contact matrix 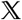 we estimate library size *L* by the sum of the upper triangular matrix of each of the chromosome specific contact matrices. This discards inter-chromosomal contacts as well as the diagonal of the contact matrix. The logCPM matrix 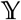 is defined as

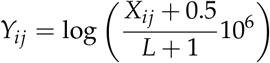

where *X*_*ij*_ refers to element *i*, *j* from the contact matrix 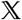 and *L* is the estimated library size for that matrix. For data normalized using HiCNorm both 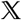 and *L* are not integers.

### ICE

For running ICE on the lymphoblastoid cell line data, we used an implementation of the algorithm as described in (Fortin, Hansen, 2015), with a tolerance of 10^−3^. We applied our implementation of ICE to unnormalized Hi-C count matrices.

For running ICE on the Greenwald data, we used the ICE implementation in Juicer (Durand et al., 2016), with the default filtering steps.

In addition, we also applied the observed-expected transform (Rao et al., 2014) to ICE-transformed data. Because the observed-expected transform removes the decay of distance, to preserve the normalization performed by the transform while still allowing the contact matrix to exhibit a distance-dependent decay, we defined a backsolve operation. For each matrix band, we compute a mean band by first computing the mean of a given band for each sample, and then the mean of these means. This latter quantity is our band mean. We then multiply each element in each sample for a given band by this mean band value. In this way we allow for the inter-sample normalization to be preserved while re-introducing a distance-based decay.

### HiCNorm

We use HiCNorm (Hu et al., 2012) in Figure 3. To process the data we used an updated implementation (https://github.com/ren-lab/HiCNorm). Following HiCNorm normalization, we applied the log counts per million transformation (see above). We then smoothed the contact matrices with a box smoother with a bandwidth of 5 bins; we use HiCRep to choose the bandwidth based on the correlation between technical replicates (T Yang et al., 2017). The bandwidth we select is the same as the bandwidth selected for 40kb resolution Hi-C data in T Yang et al. (2017). Smoothing was performed using the EBImage package (Pau et al., 2010); this is a separate but equivalent implementation to Hi-CRep.

### BNBC

BNBC has the following components: separate smoothing of each contact matrix, application of the band transformation, quantile normalization on each band matrix and finally application of ComBat on each band matrix.

Following the log counts per million transformation of the raw contact matrices, we smooth individual chromosome matrices using a box smoother with a bandwidth of 5, as selected by the HiCRep approach (T Yang et al., 2017). Each contact matrix and each chromosome is smoothed separately. We next apply the band transformation (see above) and quantile normalize each band matrix separately (Bolstad et al., 2003). Smoothing and quantile normalization is optional in our implementation; our experience suggests the these two steps have negligible impact on the performance of removing unwanted variation (particularly with respect to *R*^2^), as long as the rest of BNBC is performed. Smoothing does increase the between-sample correlation, as reported by HiCRep (T Yang et al., 2017).

Following quantile normalization we apply ComBat (Johnson et al., 2007) to each band matrix separately. We apply the parametric prior described in Johnson et al. (2007). Prior to applying ComBat, we filter out matrix cells for which the intra-batch variance is zero for all batches. After applying ComBat we set filtered matrix cells to zero. Using ComBat with a batch factor is a variant of regressing out the batch factor for each contact cell, using an Empirical Bayes approach to improve power in small sample situations as well as allowing for variances to differ across the level of the batch factor.

In our application of BNBC to data from chromosome 22, the 8 most distant bands (corresponding to loci separated by 88 Mb) were set to zero to avoid fitting ComBat to very sparse data.

Our implementation of BNBC is available in the bnbc R package from the Bioconductor project (Gentleman et al., 2004; Huber et al., 2015) at https://www.bioconductor.org/packages/bnbc.

### Explained variation and smoothed boxplot

To assess unwanted variation for each matrix cell in a contact matrix, we employ a linear mixed model approach. Specifically, we fit a mixed effect model regressing HiC contact strength on batch indicator, with a random effect at the subject level to capture the increased correlation between technical replicates. This model is fit using the R package *varComp* (Qu et al., 2013) and *R*^2^ for this model is calculated using the method of Edwards et al. (2008). The model takes the form

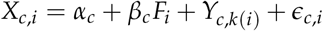

where *X*_*c*,*i*_ is matrix cell *c*, sample *i* and *α*_*c*_ is an overall scalar mean parameter. *β*_*c*_ is a vector with 3 entries (assuming 3 batch levels), constrained to sum to 0 and *F*_*i*_ is a factor vector (ie. a vector with 3 elements, 2 of which are zero, the last is 1 describing which batch a given sample comes from). 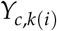 is a Gaussian random effect (ie. it has zero mean and a variance which models how correlated two replicates from the same individual are) with *k*(*i*) denoting which individual sample *i* comes from and finally *ϵ*_*c,i*_ is a symmetric, zero-mean, error term.

To display *R*^2^ as a function of distance, we first compute a series of box plots of *R*^2^, one for each band matrix.

We extract the summary measures for the box plots (median, 1st and 3rd quantile and 1.5 times the inter-quartile range). We then display these 5 curves, with color fills. Medians are black, 1st and 3rd quartiles are pink and 1.5 times the inter-quartile range are blue.

### A/B compartments from smoothed contact matrices

A/B compartments were originally proposed to be estimated using the first eigenvector of a suitable transform of the contact matrix Lieberman-Aiden et al., 2009. Specifically, the contact matrix was transformed using the observed-expected transformation where each matrix band was divided by its mean. Our contact matrices following application of the log counts per million transform and smoothing are on the log scale. To get A/B compartments from the output of BNBC (Supplementary Figure 7), we exponentiate every entry in the matrix, multiply by 10^6^, apply the observed-expected transformation and compute the first eigenvector. Data are then smoothed using a 3 bin moving-average as done by Fortin, Hansen (2015).

### QTL Study

To assess the downstream impact of the different possible normalization schemes, we conducted a study to find genetic variants associated with quantitative Hi-C signal in a given contact matrix cell when observed across 9 replicate-level observations from 5 unrelated individuals (Table 1); we refer to these variants as quantitative trait loci (QTLs). Genotypes were obtained from 1000 genomes (1000 Genomes Project Consortium et al., 2015); a detailed description is available in Gorkin et al. (2019).

As candidate SNPs we consider SNPs for which at least 2 genotypes (i.e. from a variant with alleles *A* and *B*, out of 3 possible genotypes AA, AB, and BB, at least 2 are observed) and each observed genotype has at least two subjects represented (i.e. if AA and AB are observed, at least 2 subjects have the AA genotype and at least 2 subjects have the AB genotype). Furthermore, a candidate SNP for a given contact cell is required to sit in one of the two anchor bins of the contact cell.

For a given Hi-C contact matrix cell, we specifically model the observations of this contact matrix cell, over all 9 replicates from all 5 individuals, using a mixed effect model to account for subject-level correlation in the replicate-level observations. We model the impact of genotype as a fixed dosage effect. We include as covariates the reported ethnicity of each subject (Table 1), as well as the first 3 genetic PCs, computed using SNPRelate (Zheng et al., 2012). P-values were computed by Wald test on the fixed effect coefficient for genotype, with degrees of freedom estimated via Satterthwaite’s method, as implmented in Kuznetsova et al. (2015) and Bates et al. (2015).

We conducted this study using all Hi-C matrix cells for chromosome 22 for all Hi-C matrix bins separated by no more than 700 40kb bins (2.8e7bp). We required each variant we tested to have in-sample at least 2 unique genotypes and at least 2 observations in at least 2 unique genotypes. These criteria resulted in 1,111,408 tests involving 22,593 unique SNPs and 872 unique 40kb bins on chr22.

### Greenwald data access

All data from Greenwald et al. (2019) was obtained from GEO URL: https://www.ncbi.nlm.nih.gov/geo/query/acc.cgi?acc=GSE125540. Information on batch, processing date and experimental design kindly provided by Kelly Frazer and Anthony Schmitt. Data were extracted from .hic files using Straw (Durand et al., 2016) and further read into R.

### Greenwald batch variable definition

Examining sample library sizes from chr22 (see 3) indicated a strong bifurcation of samples into those with either *>*1,000,000 reads or *<*1,000,000 reads. Upon communication with Drs. Frazer and Schmitt, it was discovered that multiple samples were combined into one .hic file, for those subjects for whom multiple observations were taken in one cell type. These individuals are the same ones with *>*1,000,000 reads in chr22 matrices, and the column **1 Rep** indicates whether a sample is coming from one replicate (*True*) or more than one replicate (*False*).

To correct for both this increased library size (above and beyond what is accounted for in logCPM transform, see below) and the effect of experimental batch, we define a new variable, **replicate-batch**, which groups samples within a batch into 2 groups, those with only one replicate or those with more than one replicate. This new variable **replicate-batch** is used for analyses of batch effect and batch effect correction.

### Sparse matrix library size correction and normalization

To accommodate the inherent sparsity of 5kb resolution Hi-C data, we employ a modified logCPM transformation as recommended by A Lun (2018):

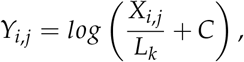

where *i*, *j* index a matrix cell, *X*_*i,j*_ is the integer count value for cell *i*, *j*, and *k* indexes samples such that *L*_*k*_ is the library size (defined to be the sum of the upper triangle for each chromosome separately) for sample *k*, *k* ∈ 1 … *n*, and

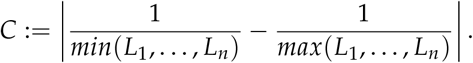

Following application of this modified logCPM to each sample’s chr22 contact matrix, we applied smoothing with a window of 5 bins (which amounts to averaging over a local 25kb neighboarhood). After dropping one sample due it being a singleton **replicate-batch**, we then applied BNBC to the first 200 matrix off-diagonals, protecting for the effect of cell type in the process of normalization.

### Evaluation of batch effect conditional on cell type

To measure the unwanted variation attributable to batch while conditioning away variation attributable to cell type, we use the coefficient of partial determination, or partial *R*^2^:

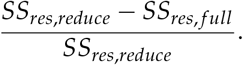

Here *SS*_*res,reduce*_ is the sum of squared residuals from the reduced model that has only cell type, and *SS*_*res,full*_ is defined similarly, but the model contains also *replicate-batch*.

### Differential Contact Analysis

To characterize the discovery power of BNBC, we performed a differential contact analysis, both on data that has been transformed by ICE-OE, and also with BNBC applied. We run ICE followed by OE on one sample’s contact matrix at a time as above, although we use the ICE implementation of HiCBricks (Pal et al., 2019) for speed. For each contact matrix cell in the first 200 matrix bands, we test for significant differences between the 2 cell types. To assess statistical significance, we use IHW (Ignatiadis et al., 2016) to model FDR conditional on the distance between the interacting loci, with a nominal *α* = 0.05. Finally, we evaluate ‘discoveries’ at IHW-FDR thresholds of 0.05, 0.01 and 0.001, as well as a Bonferroni threshold defined to be 0.05 divided by the number of tests possible.

## Data Availability

Compartments (Figure 7), QTL results (Figure 8) and differential enrichment results (Figure 10) are available from figshare, DOI: doi:10.6084/m9.figshare.c.5254002.

## Disclaimer

The content is solely the responsibility of the authors and does not necessarily represent the official views of the National Institutes of Health.

